# Maternal and zygotic factors sequentially shape the tissue regionalization of chromatin landscapes in early vertebrate embryos

**DOI:** 10.1101/2021.04.14.439777

**Authors:** Kitt D. Paraiso, Ira L. Blitz, Ken W.Y. Cho

## Abstract

One of the first steps in cellular differentiation of vertebrate embryos is the formation of the three germ layers. Maternal pioneer transcription factors (TFs) bind to the regulatory regions of the embryonic genome prior to zygotic genome activation and initiate germ layer specification. While the involvement of maternal TFs in establishing epigenetic marks in whole embryos was addressed previously, how early pluripotent cells acquire spatially restricted epigenetic identity in embryos remain unknown. Here, we report that the H3K4me1 enhancer mark in each germ layer becomes distinct in germ layer specific regulatory regions, forming super-enhancers (SEs), by early gastrula stage. Distinct SEs are established in these germ layers near robustly regulated germ layer identity genes, suggesting that SEs are important for the canalization of development. Establishment of these enhancers requires a sequential function of maternal and zygotic TFs. By knocking down the expression of a critical set of maternal endodermal TFs, an overwhelming majority of the endodermal H3K4me1 marks are lost. Interestingly, this disappearance of endodermal marking coincides with the appearance of ectodermal and mesodermal H3K4me1 marks in the endoderm, suggesting a transformation in the chromatin state of these nuclei towards a more ecto-mesodermal state. *De novo* motif analysis to identify TFs responsible for the transformation recovers a profile for endodermal maternal TFs as well as their downstream target TFs. We demonstrate the importance of coordinated activities of maternal and zygotic TFs in defining a spatially resolved dynamic process of chromatin state establishment.

## Introduction

Following fertilization, the newly formed embryo (zygote) transitions from a pluripotent state to a collection of multipotent zygotic cells. During this process the transcriptionally silent embryonic genome undergoes zygotic genome activation (ZGA), to initiate the establishment of developmental programs for the three primary germ layers. Crucial to this process is the action of maternal products, including transcription factors (TFs), which are loaded into the egg during oogenesis. While the role of maternal TFs in coordinating the initial transcription of the zygotic genome and germ layer differentiation has been extensively studied (1, 2), their involvement in the differentiation of chromatin landscape is poorly understood. Examining the role of TFs on chromatin in developing embryos is challenging due to a rapid rise in cellular heterogeneity in the embryo, along with morphogenetic movements affecting cell-cell interactions. However, by focusing on the role of maternal TFs in sculpting the chromatin landscape during the earliest stages of germ layer specification, when transcription from the embryonic genome has not yet begun and the number of different cell types is minimal, we can avoid these complications.

In *Xenopus* embryos, the three germ layers form along the animal-vegetal (AV) axis. Specifically, ectoderm is formed in the animal pole, endoderm is formed in the vegetal pole, and mesoderm is induced in the equator between the animal and the vegetal poles. During the early cleavage divisions maternally deposited mRNAs and proteins are inherited by individual blastomeres. Many maternal products show localized expression, thereby giving rise to an asymmetric segregation of cytoplasmic factors, including TFs, which affect cellular differentiation (3). By altering transcriptomic profiles of cells along this initial AV axis and rapidly reprogramming the chromatin state of the embryonic genome, maternal TFs drive differential specification of germ layer cell identities.

After fertilization, the vertebrate genome remains naïve, unmarked epigenetically and undifferentiated for a limited period of time (4–7). Recent work has shown that a network of maternal TFs encoding Fox, Sox and Pou family proteins acts through conserved mechanisms to reprogram the cellular genome into the early embryonic states (8–13). Our recent work examining the function of vegetally localized maternal Otx1 and Vegt, together with Foxh1 show combinatorial binding of these TFs forming enhanceosome complexes on the enhancers of target genes, that later become docking sites for numerous zygotically-active TFs for endoderm specification (14). These enhancer marks are established during early germ layer specification and we and others showed that their highest signal occurs on cis-regulatory modules (CRMs) that are clustered, forming super-enhancers (SEs) in the vicinity of developmentally relevant genes (11, 14). These SEs mark key cell identity genes and activate reporter expression more strongly (15), although their function within an organism is poorly understood.

In the present study, we examined the contribution of maternal and zygotic factors for the chromatin landscape changes surrounding germ layer specific enhancers and SEs. We find that distinct germ layer-specific SE profiles are frequently mapped near key germ layer patterning genes, and established by early gastrula. Interestingly, these SE associated genes are more robustly expressed compared to other genes in the genome, suggesting that SEs are important for developmental canalization. Inhibition of zygotic gene expression using α-amanitin and depletion of endodermally localized maternal TFs show that the spatially restricted SE marking along the AV axis requires the inputs of both maternal and zygotic TFs, and that in the absence of maternally deposited endodermal TFs, the endodermal chromatin state in vegetal tissue transforms towards a more mesoderm/ectoderm state. This work demonstrates the importance of coordinated activities of maternal and zygotic TFs in defining a regionally resolved dynamic process of chromatin modification in the early embryo.

## Results

### Establishment of distinct chromatin state during gastrula stage

To understand the establishment of tissue-specific chromatin states, we dissected fixed *Xenopus* gastrulae (NF 10.25, 7 hours post-fertilization or hpf) along the AV axis into the ectoderm, mesoderm and endoderm germ layers. The chromatin state of regulatory regions (i.e. promoters and enhancers) was interrogated by performing H3K4me1 ChIP-seq and compared between the different tissues. By the early gastrula, select loci surrounding developmental genes already appear distinct between the different germ layers (Figure 1A). For example, genes that are zygotically expressed in the endoderm (vegetal-most) such as *gata6, nodal* and *hnf1b* show a vegetal-animal gradation in H3K4me1 marking. Conversely, genes that are zygotically expressed in the ectoderm (animal-most) such as *bambi, foxi1* and *klf17*, show the strongest signals in the ectoderm sample, although the gradation is not as stark. The H3K4me1 markings are not limited to genes involved in germ layer patterning. We find that as early as the gastrula stage, genes involved in anterior-posterior (AP) regionalization of the embryo are already marked during the start of gastrulation. All four Hox clusters, which contain genes that regulate AP cell identity (16), are marked by H3K4me1 in all three germ layers (Figure S1) prior to their expression. The 38 annotated Hox TF encoding genes (17) are expressed poorly during this stage of development as only 3 are expressed at a transcripts per million (TPM) > 1, and 16 of which have no detectable expression (TPM = 0) (18, 19). Only when the embryos reach late gastrulation (NF 12.5, 10 hpf) do the majority of Hox genes show an expression of > 1 TPM. H3K4me1 signal in the early gastrula demonstrates pre-marking of the Hox regulatory code prior its establishment of the AP axis. By performing peak calling on these ChIP-seq datasets, the majority of H3K4me1 peaks already appear to be distinct between each germ layer (Figure 1B, S2), while the rest are shared between two or all of the germ layers (including peaks in the Hox gene cluster).

**Figure 1.**
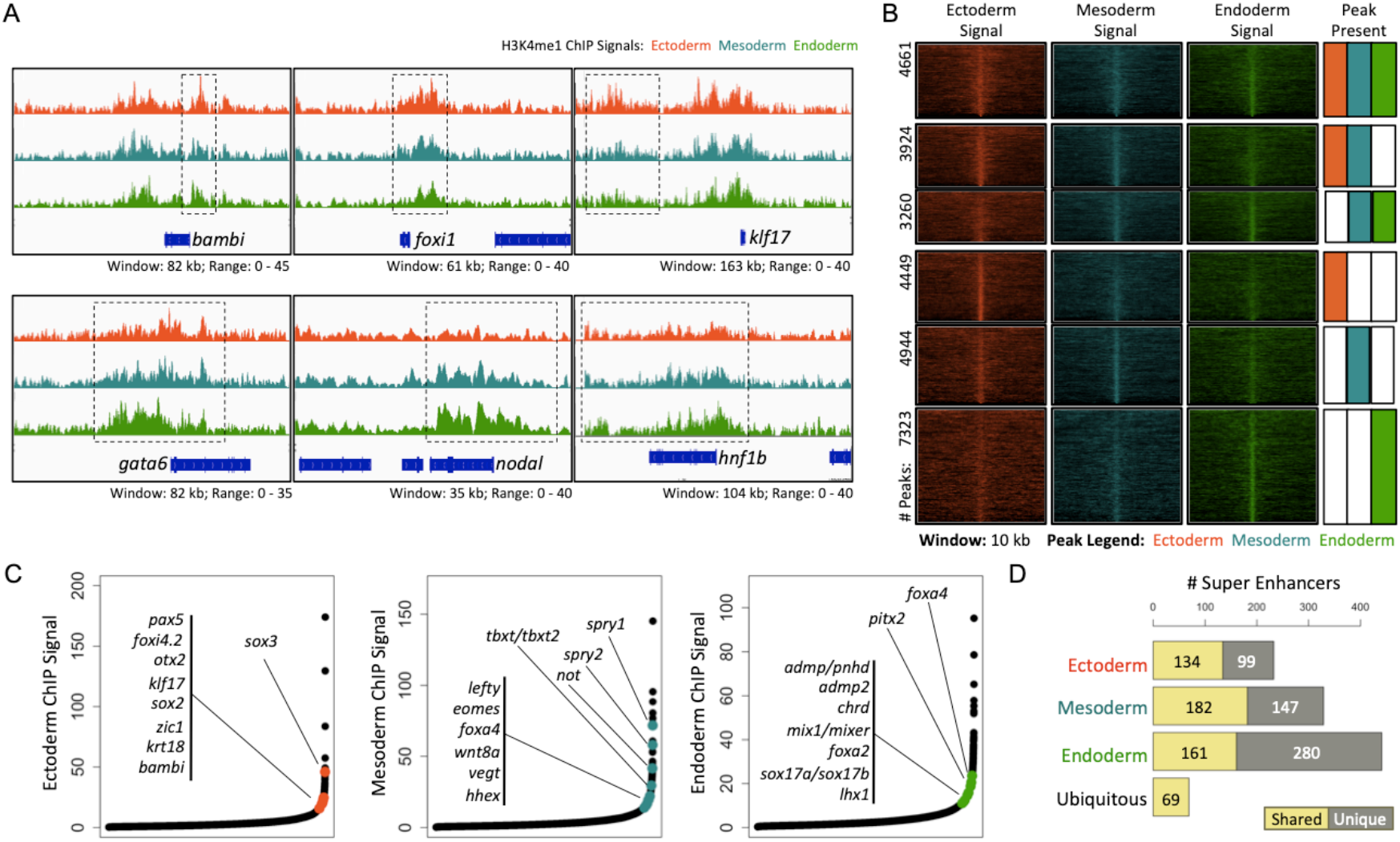
Enhancers and super-enhancers of the three germ layers are distinct by early gastrula. (A) Genome browser of H3K4me1 ChIP-seq obtained from dissected ectoderm, mesoderm and endoderm near developmental genes. (B) Germ layer-specific H3K4me1 ChIP-seq signal in different combinations of ectoderm, mesoderm, and endoderm peaks. (C) Ranking of H3K4me1 signal in enhancer regions to identify SEs. In black are all identified enhancers obtained from the H3K4me1 ChIP. Highlighted are genes known to be important for each germ layer that is associated with a SE. (D) Overlap of SEs domains from the three germ layers showing germ-layer specific and germ-layer shared SEs.

Previously, we showed that the vegetal H3K4me1 marking can be used to identify endodermal super-enhancers (SEs), which are enriched near endodermally expressed genes (14). To further study the difference of between the germ layer chromatin states, we examined H3K4me1 signal ranking to obtain a set of SEs of each germ layer (Figure 1C).

Manual inspection of SEs for each germ layer shows markings near genes that encode for regulators and markers representative of each tissue. Ectodermal SEs are located near genes that encode ectodermal TFs such as *sox3* (20), and epidermally-expressed cement gland cytokeratin *krt18* (21). In the mesoderm, SEs mark genes encoding T-box TFs *tbxt* (22), *eomes* (23) and the zygotically-expressed *vegt* (24–27); as well as negative feedback regulators of Fgf signaling *spry1* and *spry2*, which regulate the morphogenetic movements of mesodermal cells (28). Endoderm SEs mark critical regulators of endodermal fate including *foxa4* (29), *mix1* (30), *mixer* (31), *sox17* (32), and *lhx1* (33). These results suggest that many TFs known to be critical for germ layer specific properties are closely associated with the germ layer specific SEs (Figure 1D). We conclude that the H3K4me1 marked enhancers are already regionally distinct along the AV axis by the early gastrula stage.

### Tissue-specific SE marking is associated with spatial restriction and robustness in gene expression

While the role of promoters and enhancers during development has been well characterized, the role of SEs in developmental systems is yet unclear. Previously, we identified endodermal SEs near genes coding for key regulators of endodermal fate that are activated during ZGA (14). Here we examine the formation of germ layer-specific SEs during spatial patterning of embryos. We assigned each SE to the nearest gene(s) (Figure 2A), and examined whether the transcriptional activity of the genes associated with SEs are germ layer specific and quantitatively different from that of the other genes that are not associated with SEs. While many SE-associated genes showed specificity for each germ layer (92 for ectoderm, 144 for mesoderm, and 281 for endoderm), 95 SEs are H3K4me1-marked across all three germ layers. This set of genes includes developmental genes that are required for tissue regionalization across both the dorsal-ventral (DV) axis (e.g. *ventx1*.*1* and *ventx2*.*1*) and the AP axis (e.g. *hoxb2* and *hoxb4*) rather than germ layer formation along the AV axis. SE-associated genes are more highly expressed than non-SE associated genes (15), which we consistently find in our early embryonic tissues for all three germ layers (Figure 2B). Further, the expression levels of each set of SE associated genes show distinct spatial pattern consistent with their germ layer-specific H3K4me1 marking (Figure 2C). Ectoderm and endoderm SE-associated genes are expressed in animal-vegetal and vegetal-animal gradients, respectively. Mesoderm SE-associated genes are expressed in the embryo equator, and less so in the poles.

**Figure 2.**
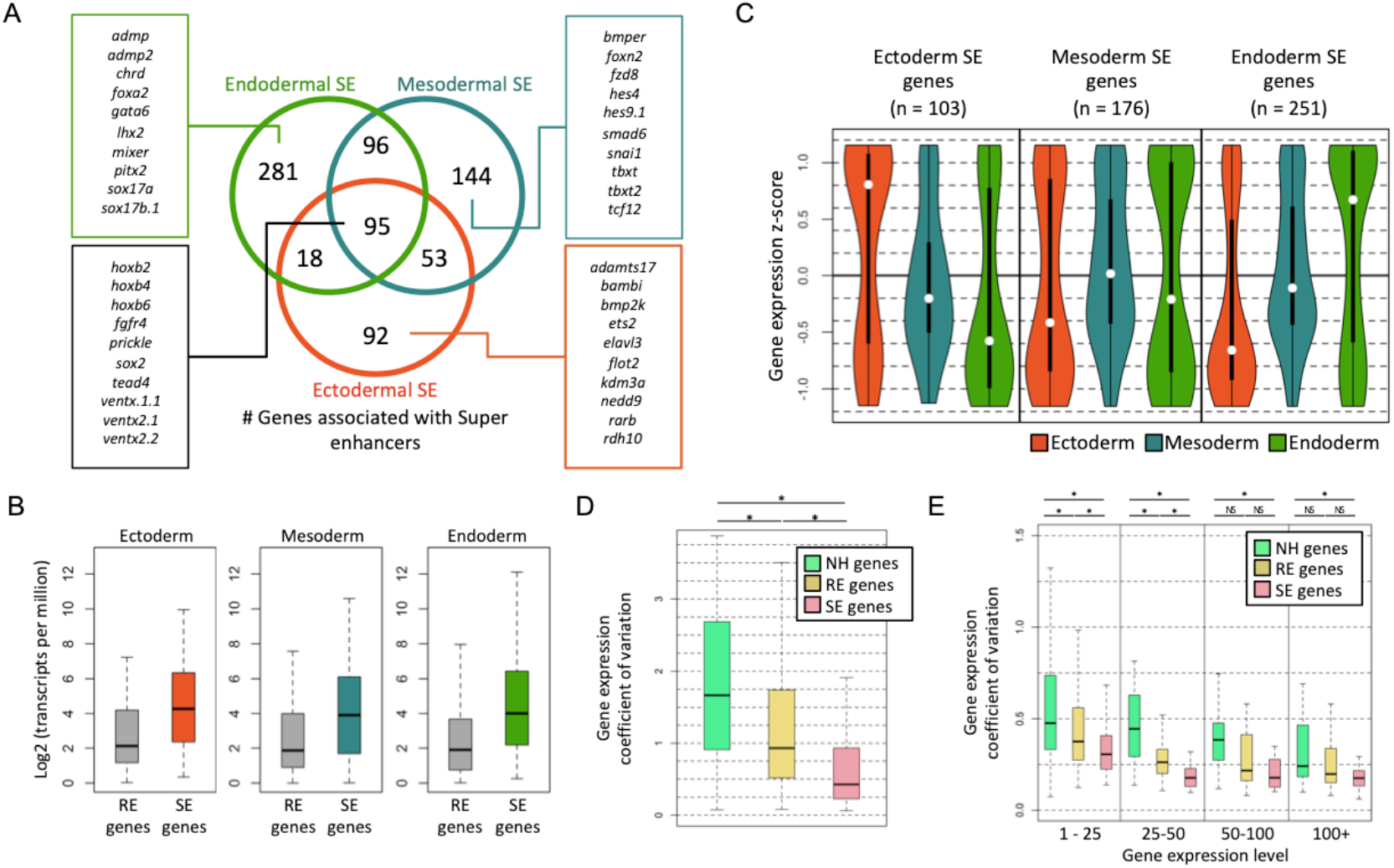
Genes associated with super-enhancers are tissue-specific and robustly expressed during development. (A) Genes associated with SEs for the three germ layers. (B) Expression of genes associated with either regular enhancers (REs) or SEs in their respective tissues. (C) Distribution of the expression of genes associated with SEs and their expression in the ectoderm, mesoderm and endoderm. (D) Coefficient of variation (COV) of gene expression from the RNA-seq of 15 embryos from 3 clutches at 7 hpf. Displayed are the COV for non-H3K4me1 (NH) associated, RE- and SE-associated genes. (E) COV of gene expression for genes that are either non-H3K4me1 (NH) associated, RE- or SE-associated genes. The expression levels are binned as 1-25, 25-50, 50-100, and 100+ TPM to account for variation caused by the noisy expression at low TPMs. Significance is tested by Wilcoxon Rank-Sum test. * = p-value < 0.01 and NS = not significant.

SEs have been proposed to act in a combinatorial fashion in gene regulation to confer robustness in gene expression (34–36). We therefore tested whether SE associated genes are expressed with lower expression variability than other genes. Multiple time-course transcriptomic datasets (collected at 30 minutes interval) from different embryo clutches were used (18, 37; Figure S3A) to compare the coefficient of variation (COV) of zygotic gene expression in transcripts per million (TPM) across 11 time points from 4 hpf (NF 8) to 9 hpf (NF 11.5-12). Both SE- and regular enhancer (RE)-associated genes showed lower variation than genes that are non-H3K4me1 (NH) associated (Figure S3B). Because RNA-seq is generally noisy at lower levels of gene expression, we binned gene expression based on TPM levels and compared COV of genes in associated bins. In the lowest TPM bin, genes show relatively similar COV across all three sets (Figure S3C). At higher TPM bins, where there is less technical noise, the NH genes maintain a similar level of COV as those in the low TPM bin. Genes associated with REs have lower COV than NH genes. Even more so, genes associated with SEs showed the least variation.

Since these RNA-seq datasets are generated from pooled embryos, we also measured the transcriptome of 15 individual embryos from 3 different clutches (5 each) at 7 hpf (NF 10.25) so we can measure the COV at the individual embryo level (Figure S3D). Embryos within the same clutch showed more similar transcriptomes (Figure S3E). For each gene, we measured the COV. Among the genes with the least variation are well known germ layer patterning genes (e.g. *crx, sox17a, tbxt* and *foxh1*.*2*) and dorsal-ventral patterning genes (e.g. *ventx2*.*2, wnt5b, fgf8* and *lefty*), many of which encode signaling molecules and transcription factors. Among the top 100, 37 are associated with SEs and another 37 are associated with REs, suggesting enrichment for genes with H3K4me1 marks. Indeed, SE associated genes are enriched in low COV in expression (Figure S3F). As a group, SE associated genes tend to have the lowest COV, followed by RE associated genes (Figure 2D). When comparing different expression level bins, SE associated genes have the lowest variations in expression (Figure 2E), consistent with our previous findings.

Overall, our analysis shows two putative regulatory functions of SEs during embryo development. Early established SEs are associated with both spatial patterning and also robust expression of genes.

### Maternal and zygotic requirements for establishment of chromatin states

Previous work has shown that maternal TFs play critical roles in the establishment of chromatin states using bulk embryo samples (8, 10-14, 38, 39). However, how maternal TFs contribute to the establishment of germ layer-specific SEs is unclear. To delineate the roles of maternal and zygotic factors in the early stages of chromatin differentiation, a RNA polymerase II inhibitor, α-amanitin, was microinjected into 1-cell stage embryos (40). We dissected early gastrula (NF 10.25, 7hpf) embryos with and without α-amanitin into the three germ layer regions and compared H3K4me1 marks. The loss of zygotic factors exerted variable effects on H3K4me1 deposition in different peak regions (Figure 3A). Near genes such as *not*, the H3K4me1 marking appeared largely unchanged across all three germ layers when comparing α-amanitin to uninjected embryos, suggesting a predominantly maternal control of H3K4me1-marking for these regions. In the loci surrounding genes which are expressed in an animally-enriched manner such as *bambi* and *klf17*, the H3K4me1 marking is largely unchanged in the ectodermal and mesodermal tissues.

**Figure 3.**
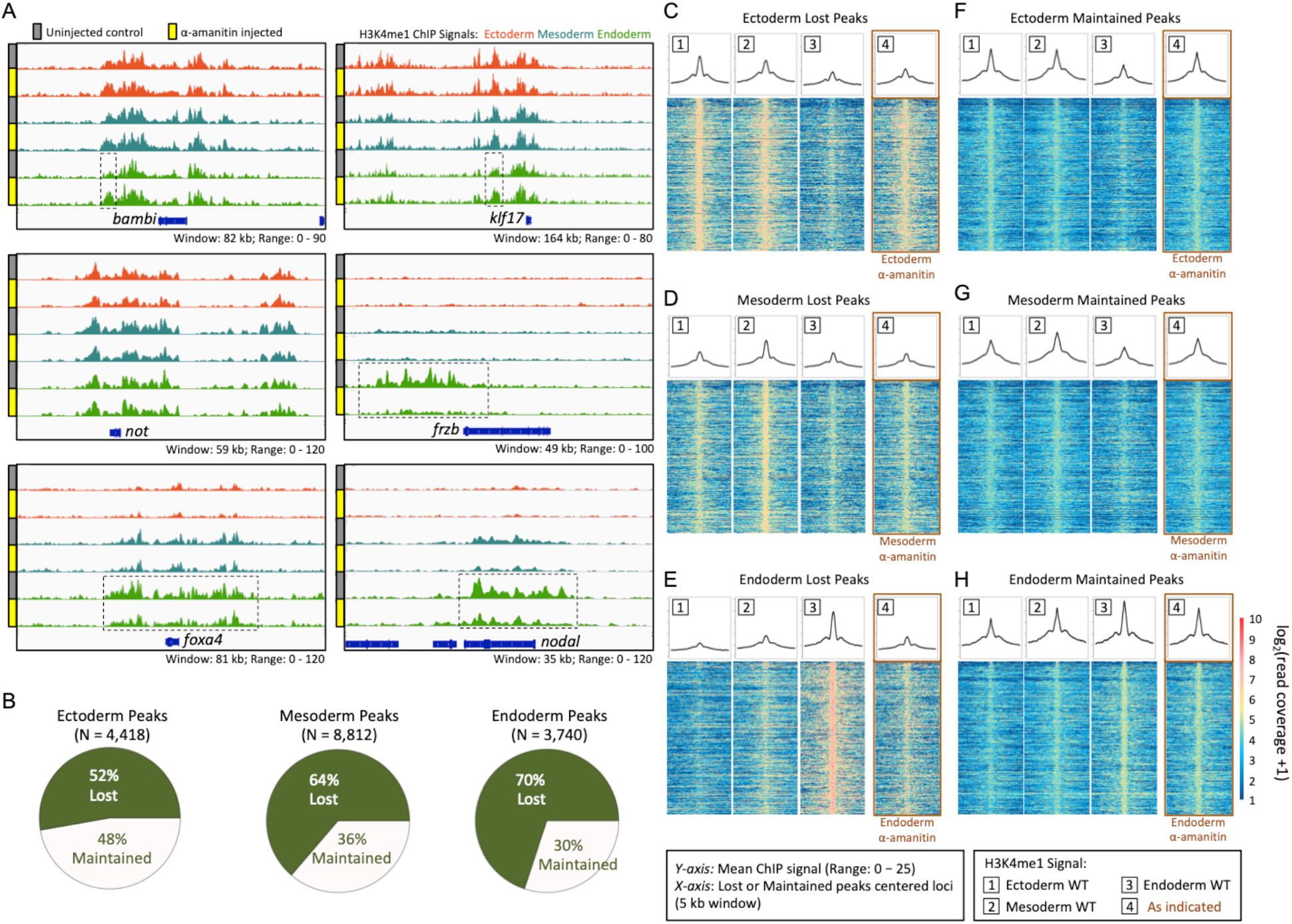
Maternal and zygotic requirements for establishment of cis-regulatory landscape. (A) Genome browser of H3K4me1 ChIP-seq obtained from germ layer tissues of control and α-amanitin injected embryos in the *not, klf17, bambi, foxa4, frzb*, and *nodal* loci. (B) Fraction of lost and maintained wild type ChIP-seq peaks after α-amanitin treatment. (C-H) The heatmap shows read coverage of the wild type H3K4me1 signal in the three germ layers centered on the peaks with a 5 k.b. window that were either ‘lost’ (C-E) or ‘maintained’ (F-H) after α-amanitin injection. The line plot above each heatmap is a measure of the mean ChIP-seq signal for each base within the 5 k.b. window.

However, these marks are somewhat increased in the endodermal tissue in the α-amanitin injected samples, particularly the marks upstream of the gene body, suggesting that the establishment of H3K4me1 marks in these regions is inhibited by zygotic factors in the endoderm. In the meso-endodermally- or endodermally-expressed genes (e.g. *foxa4, frzb* and *nodal*), H3K4me1 is more prominent in the endoderm, showing a distinct vegetal-animal gradation in uninjected embryo tissues. The marking of these genes in the endoderm is severely disrupted in the presence of α-amanitin, and resembles that of the wild type ectodermal or mesodermal tissue. The reduction in H3K4me1 marking spans promoter-proximal, intronic and intergenic regions suggesting that the deposition of both promoter and enhancer markings are inhibited. This result suggests that zygotic factors are required for the deposition of H3K4me1 marking in these regions.

To quantify the effects of maternal and zygotic factors in establishment of these marks, we performed peak calling on the H3K4me1 ChIP-seq datasets in both control and α-amanitin samples. We counted the number of peaks that were either maintained or lost after α-amanitin treatment. The maintained peaks represent maternally controlled peaks, whereas lost peaks in the presence α-amanitin represent zygotically controlled H3K4me1 modification. Previous bulk embryo chromatin datasets have suggested varying levels of maternal control of chromatin states. For example, H3K4me3-marked promoters and H3K27me3-marked repressed regions are largely maternally-controlled such that α-amanitin treatment resulted in only 14% and 10% of peaks lost, respectively (7). Ep300-marked enhancers on the other hand lost 85% of peaks after α-amanitin treatment, suggesting strong dependence on zygotic factors for the binding of this enhancer-marking factor (7). We find that H3K4me1 marking appears to be intermediate between these two extremes (Figure 3B). The percentages of peaks dependent on zygotic factors varies between the ectoderm, mesoderm and endoderm - 52%, 64% and 70% of peaks were lost, respectively, upon α-amanitin treatment. This suggests that maternal and zygotic factors collaborate to control the early chromatin states. In addition, the balance of control between maternal and zygotic factors varies between the three germ layers in establishing the early epigenome.

To better understand which regions are preferentially regulated by zygotic factors, we quantified the read coverage in peaks that were lost in genomic landmarks after α-amanitin treatment. First, we compared the promoter and enhancer regions (Figure S4). Both promoter and enhancer H3K4me1 markings are disrupted by blocking zygotic transcription. H3K4me1 peaks are reduced more in the promoters than enhancers in both the ectoderm and mesoderm tissues, while the opposite is true in the endodermal tissue (Figure S4D). Second, we compared the marking in REs and SEs (Figure S5). The markings in ectodermal and mesodermal tissues are affected relatively similarly in both the RE and SE regions, whereas the endodermal tissues appear to be more affected in the SE regions (such as the SEs in the *foxa4* and *frzb* in Figure 3A). Finally, we wished to know whether the peaks that are lost (or maintained) uniformly appear across all three germ layer tissues, or whether these marks are uniquely marked in their respective germ layer. By plotting the read coverage (heatmaps) and the mean read coverage (line plot) across these lost or maintained peaks within a 5 kb span, we uncovered that lost (Figure 3C-E) and maintained (Figure 3F-H) peaks have distinct regional marking patterns. Peaks that are lost tend to have very specific marking in their respective germ layers. For example, peaks that are lost in the endoderm (Figure 3E, compare track 3 to 4) are strongly endodermally-marked showing a vegetal-animal gradation (Figure 3E, decreasing signal across tracks 3, 2 and 1). Similarly, lost mesodermal peaks (Figure 3D, compare track 2 to 4) tend to be most strongly H3K4me1 marked in the marginal region (Figure 3D, compare track 2 to 1 and 3). Finally, lost ectodermal peaks (Figure 3C, compare track 1 to 4) tend to be most strongly marked in the ectoderm with an animal-vegetal gradation (Figure 3C, decreasing signal across tracks 1, 2 and 3). These results are consistent with the view that H3K4me1 marks lost in specific germ layers are regulated by zygotically regulated TFs. Conversely, the maternally-controlled H3K4me1 marks (maintained peaks in the presence of α-amanitin) show enriched marking in their respective germ layers, and differences are much less stark (Figure 3F-H). Based on these findings, we suggest that the factors responsible for germ layer specific H3K4me1 marking are zygotic factors expressed in each germ layer. On the other hand, H3K4me1 patterns established by maternal TFs are less germ layer specific, indicating that maternally present ubiquitously expressed TFs may be responsible for majority of these histone markings. This is consistent with the Waddington landscape model of cell differentiation (41): the set of cis-regulatory domains we observe are established by sequential factors, maternal then zygotic, in different germ layers and gradually they become more divergent.

### Loss of maternal TFs results in animalization of the endodermal chromatin state

Previously we have noted the frequent correlation between the binding of three maternal endodermal factors Otx1, Vegt and Foxh1 (OVF) and their association with endodermal SEs (14). Here we investigated their direct role in endodermal chromatin state formation by knocking down the expression of these maternal TFs using morpholino antisense oligonucleotides (MOs). As with the α-amanitin experiment, OVF knockdown (KD) resulted in reduction of H3K4me1 marking in a variable manner across known meso-endodermal/endodermal genes as compared to explants from sibling control embryos (Figure 4A). The loci near genes such as *admp, pitx2, foxa4* and *lhx1* all showed decrease in H3K4me1 marking. More drastic decrease could be seen in the loci near genes such as *pnhd, hnf1b* and *gata6*. By quantifying peaks, we find that the large majority of peaks (88% of peaks) are lost after OVF MO knockdown (Figure 4B). This suggests a significant contribution of the maternal TFs Otx1, Vegt and Foxh1 in the establishment of the H3K4me1 state of the early endoderm.

**Figure 4.**
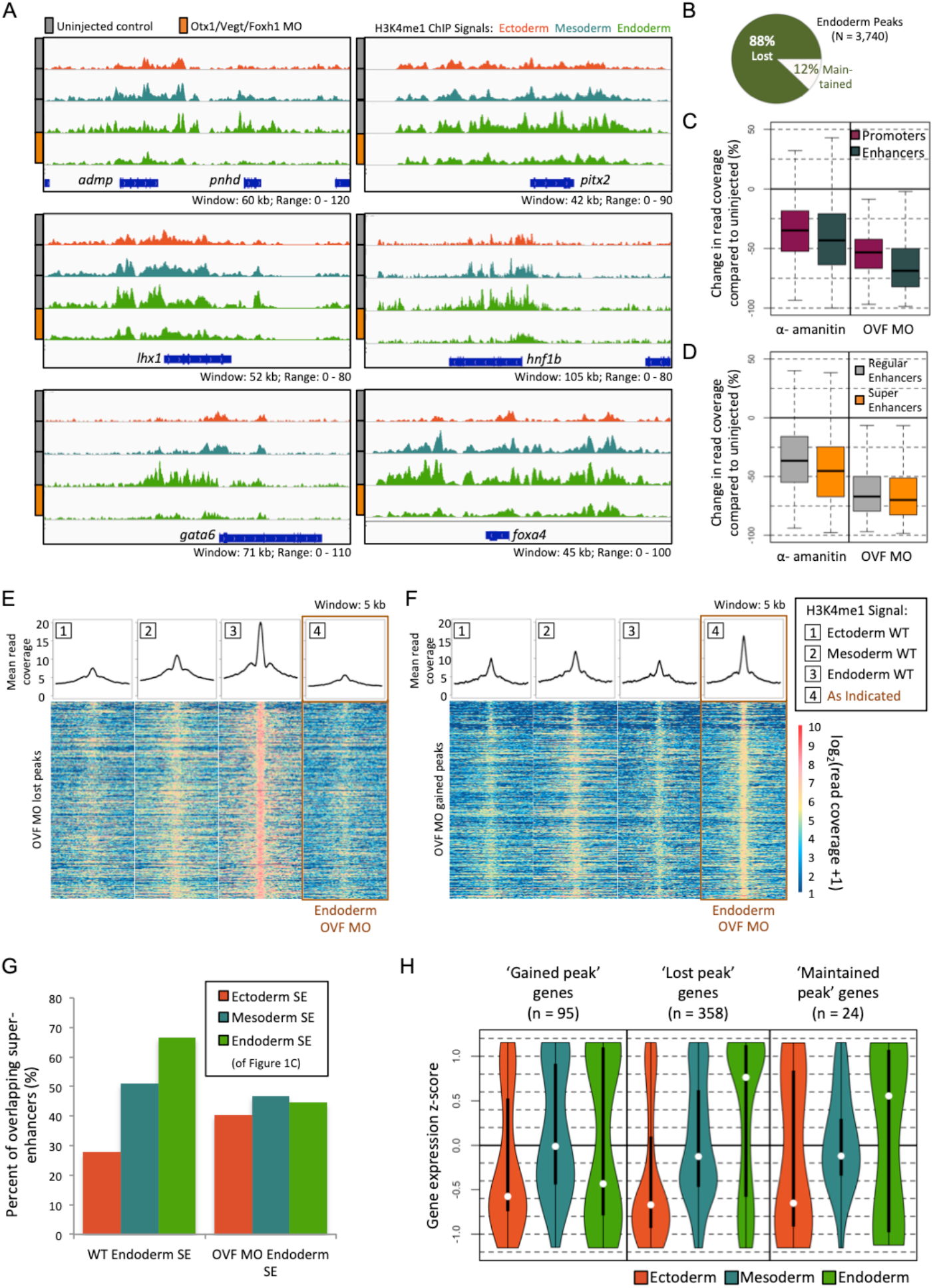
Ecto-mesodermal transformation of the endoderm chromatin state due to loss of endoderm transcription factors. (A) Genome browser of endodermal H3K4me1 ChIP-seq in control and OVF MO injected embryos. Graphed for comparison are the wild type ectodermal and mesodermal H3K4me1 ChIP-seq signal. (B) Fraction of lost and maintained wild type ChIP-seq peaks after OVF MO injection. (C) Quantification of change in read coverage in promoter and enhancer peaks after α-amanitin and OVF MO injection. (D) Quantification of change in read coverage in RE and SE peaks after α-amanitin and OVF MO injection. (E-F) The heatmap shows read coverage of the wild type H3K4me1 signal in the three germ layers centered on the peaks with a 5 k.b. window that were either ‘lost’ (E) or ‘gained’ (F) after OVF MO injection. The line plot above each heatmap is a measure of the mean ChIP-seq signal for each base within the 5 k.b. window. (G) Intersection of wild type and OVF MO injected endodermal SEs with the wild type germ layer SEs from the Figure 1C dataset. (H) Expression of genes associated with ‘gained’, ‘lost’ and ‘maintained’ peaks in the three germ layers.

To identify the genomic locations where H3K4me1 signal is lost due to the KD of OVF TFs, we first quantified read coverage in enhancers and promoters (Figure 4C, S6A). Compared to the α-amanitin treatment, the H3K4me1 marking in both promoters and enhancer regions are more significantly reduced in the OVF TFs knockdown samples. The strongest disruption is seen in the enhancers, highlighting the importance of OVF TFs in enhancer activity in the endoderm. Second, we compared the read coverage in REs and SEs. While these regions are much more disrupted in the OVF TF knockdown compared to α-amanitin treatment, effects on REs and SEs are largely indistinct (Figure 4D, S6B). Finally, we examined the regional pattern (e.g. ectoderm, mesoderm and endoderm) of the peaks that were lost or gained after OVF KD. By plotting the read coverage (heatmaps) and the mean read coverage (line plot) across these lost or gained peaks within a 5 k.b. span, we uncovered that lost and maintained peaks have distinct regional marking patterns (Figure 4E-F). Peaks that are lost (Figure 4E, compare track 4 to 3) appear to be most strongly endodermally marked with a vegetal-animal graded pattern (Figure 4E, decreasing signal across tracks 3, 2 and 1). Conversely, H3K4me1 peaks which were gained in OVF injected embryos (Figure 4F, compare track 4 to 3) tend to be most strongly marked in the mesoderm wild type sample and less so in the ectoderm and endoderm (Figure 4F, compare track 3 to 1 and 2). Further, we compared SE profiles in the control and the morpholino treated endoderm to those identified in different wild type tissues from Figure 1 (Figure 4G). We identified the coordinates of the SEs for the wild type uninjected and OVF KD and asked what percentage of these SEs overlap those found in the ectoderm, mesoderm and endoderm WT from Figure 1. The profile of the wild type unsurprisingly reflected an endodermal profile, while showing the least similarity with the ectodermal sample. On the other hand, the morpholino knockdown caused endoderm tissue to take on a SE profile that is less endodermal and more ectodermal. The match with the endoderm sample is greatly reduced, while the match with the ectodermal sample is increased.

Lastly, we associated the peaks that were gained or lost after OVF TF knockdown to the nearest neighboring gene. By plotting their expression in wild type embryos, we find that the genes that lost associated peaks were expressed in a vegetal-animal graded manner (Figure 4H). Conversely, the genes that gained peaks are expressed most strongly in the mesoderm. These patterns in gene expression mimic the patterns of H3K4me1 marking of gained and lost peaks seen in Figure 4E and F. These analyses suggest that maternal OVF TFs play roles both in establishing the endodermal chromatin state and repressing more ectodermal and mesodermal chromatin states.

### Lost H3K4me1 peaks after OVF MO are associated with the endodermal gene regulatory network

To identify a candidate set of TFs that are responsible for the modifications in H3K4me1 marking in endoderm, we performed *de novo* motif analysis on H3K4me1-flanked nucleosome-free regions (NFRs), which mark the promoter and enhancer regions of DNA that are accessible to various transcription factors. For this analysis, we utilized the Homer (42) peak calling option *–nfr*, which specifically identifies the valleys between the bimodal histone peaks. NFR peak calling of H3K4me1 signal in the control and OVF-depleted endodermal samples yielded 22,577 and 8,270 peaks, respectively (Figure 5A). Among the set of uninjected endoderm NFRs, 18,916 (84%) were lost and 3,661 (16%) were maintained after OVF MO treatment. Conversely, 4,609 NFRs were gained post-treatment.

**Figure 5.**
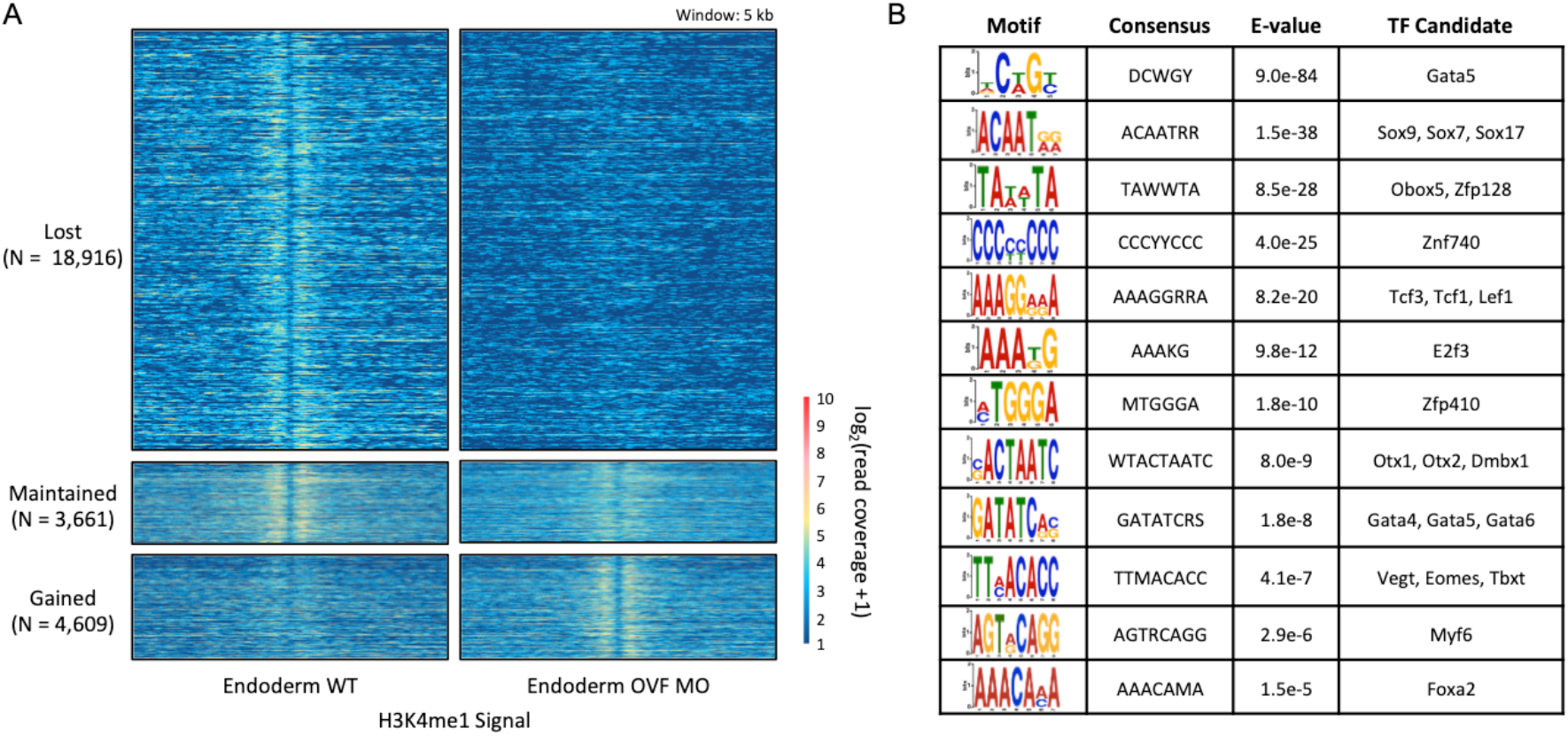
Uncovering of the endodermal gene regulatory network TF motif profile after endodermal maternal TF knockdown. (A) Heatmap of H3K4me1 ChIP-seq signal surrounding H3K4me1-flanked NFRs from wild type and OVF MO embryos. (B) Enriched motifs under H3K4me1-flanked NFRs that were ‘lost’ after maternal endodermal TF knockdown.

*De novo* motif analysis performed on the NFRs identified enrichment of both maternal and zygotic TFs consensus binding motifs (Figure 5B). The motifs identified under NFRs that were lost after Otx1, Vegt and Foxh1 KD include key TFs of the endodermal gene regulatory network. The top ranked motifs includes the T-box and paired-type homeobox motifs that are likely to represent Vegt and Otx1, respectively (14). We also found motifs representing zygotically expressed endodermal pioneer TFs Gata4/5/6 (43, 44) and Foxa2 (45, 46), as well as Sox17, which is a critical endodermal regulator (32, 47). In addition, the motif for Tcf/Lef TFs, critical mediators for patterning the DV axis of the mesendoderm, was identified (48, 49). Interestingly, we failed to identify motifs representing Smad2/3, which mediate Nodal signaling, a key endodermal signaling pathway. However, this is consistent with the finding that loss of Nodal signaling through small molecule inhibition marginally affects the H3K4me1 deposition in the early embryo (6). Conversely, the motif analysis on gained H3K4me1-flanked NFRs identified motifs of known regulators of ecto-mesodermal cell lineages (Figure S7), including the motifs for Neurog2 (50), Six1 (51) and Six6 (52, 53), Sox9 (54), Sox10 (55) and Etv2 (56). These TFs are known to regulate ecto-mesodermal derived lineages such as neurons, neural crest cells and cardiac muscle. In sum, the gained and lost H3K4me1-flanked NFR regions are enriched with motifs representing lineage specific TFs important for germ layer specification. We suggest the active involvement of these TFs in shaping the chromatin states of differentiating germ layers.

## Discussion

### Model of early chromatin state dynamics

The dynamic interplay between transcription factors, epigenetics, and cell lineage commitment during differentiation in metazoan genomes remains one of the central questions in developmental biology. Here, we generated ChIP-seq data to understand the establishment of the enhancer and super-enhancer chromatin-marking H3K4me1 within the context of cellular differentiation of the pluripotent zygote to the formation of the three germ layers. Based on our experiments using early *Xenopus* embryos, we conclude with the following model (Figure 6A). By early as gastrula stage NF 10.25 (~7 hpf, or ~2.5 hours after the initiation of the major wave of ZGA), the chromatin states of the early embryo are already distinct between the germ layers (Figure 1,2). By this point, these states appear to have been uniquely modified by both maternal and zygotic factors (Figure 3,4). These modifications appear to be intrinsically linked to the germ layer gene regulatory networks (Figure 5). Loss of endodermal maternal factors results in the transformation of the endodermal chromatin state to a more ecto/mesodermal-like state (Figure 4). The regions where histone markings are lost are enriched with the consensus binding motifs of maternal and zygotic endodermal TFs known to regulate the endodermal program (57) reinforcing the relationship between the regulators of cellular differentiation and cellular states. In the future, it would be valuable to examine the mechanism by which maternal TFs recruit histone-modifying enzymes to maintain gene regulatory programs.

**Figure 6.**
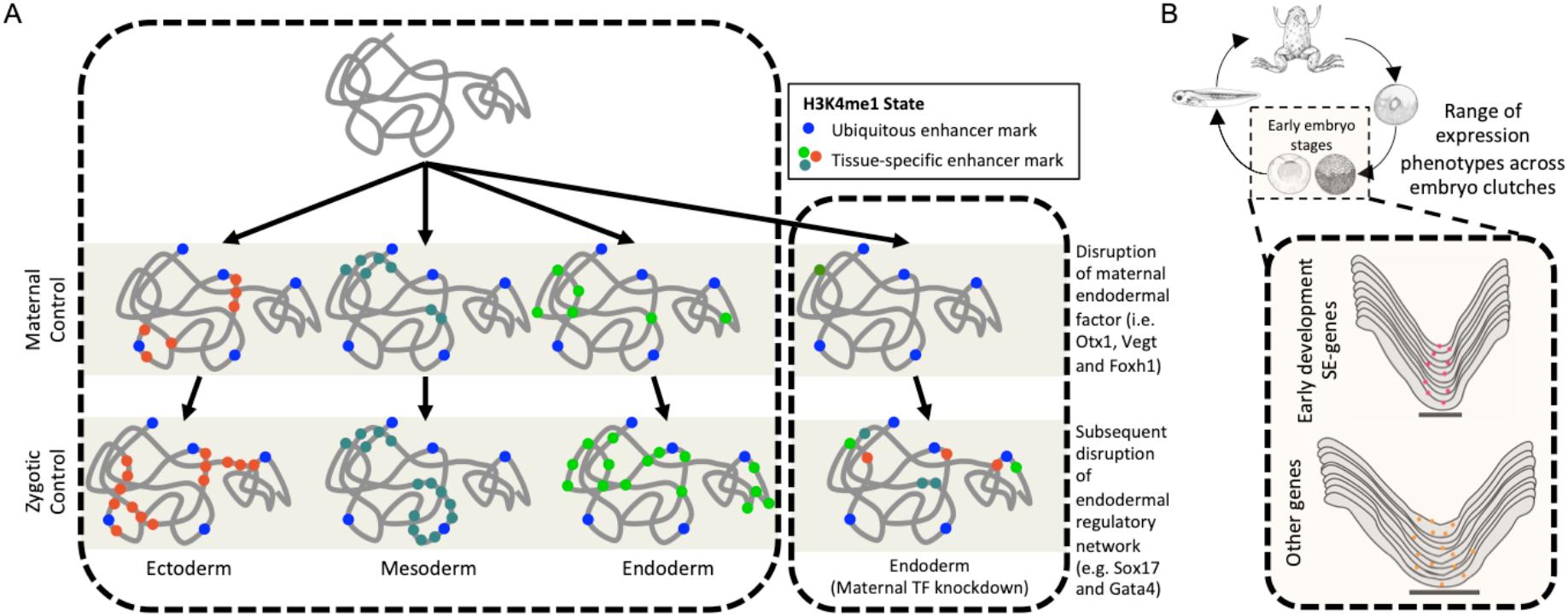
Model of H3K4me1 establishment during early embryogenesis. (A) The chromatin state differentiation during the first 7 hours of development is highly dynamic and is under both maternal and zygotic factor control. Loss of critical factors in the endoderm result in the establishment of a more ecto-mesodermal-like chromatin state in the endoderm. (B) Canalization of gene expression during early stages of development. Boxed are the early blastula and gastrula stages where gene expression and chromatin states were identified. Genes that were associated with SEs showed lower variability in expression compared to those that are non-SE associated.

### Developmental canalization, histone methylation and super-enhancers

For organisms to develop as functionally integrated systems, cellular and tissue structures have to develop in highly predictable ways. The term ‘canalization’ describes the tendency for embryonic cells or tissues to follow the same developmental trajectory during development (58). This implies that developmental mechanisms exist to drive this predictability in development. In the current study, we have examined the expression of SE associated genes by measuring variation of gene expression obtained from RNA-seq from embryos of different clutches. Since SEs are composed of multiple constituent enhancers, we reasoned that clustering of enhancers into SEs may confer robustness in gene expression, perhaps acting similar to shadow enhancers in select genes (34–36). The time course expression patterns of genes associated with SEs show that they maintain lower expression variability relative to the other genes in the genome (i.e. regular enhancer associated genes or non-H3K4me1 enhancer associated genes) (Figure 6B). Since SEs are associated with key developmental genes, they could provide buffers to variation in gene expression resulting in canalization of broad scale embryonic tissue formation. However, it remains a possibility that the histone mono-methylation itself could be the determinant of canalization of gene expression, rather than the activity of the enhancer, as has been speculated with the role of DNA methylation (59). How these H3K4me1-marked genomic elements are associated with canalization of gene expression is yet to be determined.

### Enhancers and super-enhancers roles along the anterior-posterior and dorsal-ventral axes

Our work has shown that distinct SEs are formed during germ layer specification along the animal-vegetal axis. We have in parallel identified the putative cis-regulatory regions near genes that are important for the patterning of anterior-posterior (AP) and dorsal-ventral (DV) axes. The collinearly controlled expression of homeotic (Hox) genes is important for conferring spatial identity along the AP axis (16). Among vertebrates, Hox genes are clustered on 4 different chromosomes representing *hoxa, hoxb, hoxc* and *hoxd* clusters. The majority of these genes initiate transcription during late gastrula/early neurula stage (18, 19), during which the embryo begins to extend along the AP axis. We find that all four Hox clusters are H3K4me1-marked (on both REs and SEs) by gastrula stage in all three germ layers. In embryos where zygotic transcription is inhibited, Hox clusters still remain marked with H3K4me1, suggesting that maternal factors are responsible for this early marking of these clusters. Since these loci also remain marked after the loss of the endodermal maternal OVF TFs and their downstream regulatory target genes, we propose that the H3K4me1 marking of the Hox complexes is mediated by alternative and perhaps ubiquitous maternal factors. It is unclear why the Hox genes are H3K4me1 marked hours prior to the onset of their transcription and the formation of the AP axis. Perhaps, a mechanism exists to ensure these critical genes to be protected from gene silencing during development.

Conversely, genes that are important for the DV patterning of the embryo tend to be affected by loss of zygotic transcription or loss of maternal factor function. Genes in the *ventx* cluster (*ventx1*.*1, ventx1*.*2, ventx2*.*1* and *ventx2*.*2*) (60, 61) and *bmp4* (62), which control ventral identity, are highly marked by H3K4me1 across all three germ layers. Similarly, genes such a Bmp antagonists *chrd* (63), *nog* (64) and *fst* (65), as well as Nodal ligands *nodal1* and *nodal2* (66), which are important for dorsal cell identity, are marked by H3K4me1 in the meso/endoderm germ layers. Unlike the loci near the Hox genes, these loci are partially or fully affected by the loss of zygotic transcription and therefore endodermal maternal factor function is implicated. This is consistent with the biology of the *Xenopus* embryo, whereby the orthogonal DV and germ layer patterning gene regulatory networks are entangled (57).

Little is known about the function and properties of SEs in embryos. Here, we used our ChIP-seq data to understand the role of SEs in embryos. SE studies in cell culture suggest they are particularly important for regulation of cell identity genes (15). We consistently found that SEs are important for spatially patterned gene expression during the first embryonic cell decision-making processes. Ectodermal, mesodermal and endodermal SE-associated genes are differentially expressed along the animal-vegetal axis. At present, it is unclear how far this association holds, but it would be interesting to address whether the link between SE formation and cell differentiation processes in AP and DV patterning, as well as cell differentiation at later stages similarly holds. By refining the fractionation of embryonic tissues and implementation of low input ChIP-seq-like protocols, we should be able to address these questions.

### Pioneer factors in the endodermal gene regulatory network

Pioneer factors are a class of TFs that are able to bind to closed chromatin possibly resulting in the extrusion of core histones and/or epigenetic changes important for gene regulation (46). During early development in a variety of different organisms, maternal TFs perform this role to initiate modifications to the zygotic epigenome (8, 10–14, 38, 39). In the *Xenopus* embryo, these TFs include relatives of the mammalian pluripotency TFs and germ layer-specific maternal TFs such as Foxh1, Sox3 and Pou5f3 (10, 11). In the current study, we identified a set of H3K4me1 bimodal peaks flanking nucleosome free regions (NFRs) that are lost following the knockdown of maternal endodermal TFs (Figure 5).

Consistent with the known roles of maternal TFs, these lost H3K4me1-flanked NFRs are enriched with motifs that bind maternal endodermal TFs. In addition, these NFRs are also littered with the motifs of zygotic endodermal TFs, suggesting that the set of lost H3K4me1-flanked NFRs represent the chromatin bound by the TFs in the endodermal gene regulatory cascade (57). These zygotic TFs include the *Xenopus* orthologs of the earliest mammalian pioneer TFs of hepatogenesis, the FOXA and GATA transcription factors (46), which are target genes directly regulated by Otx1, Vegt and Foxh1 (9, 14, 67, 68). Other endodermal TFs that are implicated in the process include the pan-vertebrate endodermal TF Sox17 (32). Like Foxa and Gata, the *sox17* gene is a direct target of maternal TFs mentioned above. This presents an intriguing possibility that Sox17 could also act as a pioneer factor during endoderm formation.

In sum, we have shown that maternal uniformly expressed Foxh1 together with endodermally expressed Otx1 and Vegt TFs function at the top of a hierarchy of TF interactions to not only regulate the transcriptional responses of developmental genes, but also coordinate histone modifications and affect the epigenetic landscape regulating endoderm development. Since these TFs are conserved among vertebrates, it would be important in the future to test whether the similar germ layer specific SEs are formed in different vertebrate species.

## Materials and Methods

### Animal husbandry and embryo manipulation

*Xenopus tropicalis* males and females were maintained in accordance with the University of California, Irvine Institutional Animal Care Use Committee (IACUC). *X. tropicalis* females to be used for embryo collection were injected with 10 units of Chorulon HCG (Merck and Co.) 1-3 days before and 100 units of HCG in the morning of the embryo collection. Eggs were collected in a 0.1% BSA in 1/9x MMR coated glass dish. The eggs were *in vitro* fertilized with sperm suspension in 0.1% BSA in 1/9x MMR (69) obtained from sacrificed males. The embryos were dejellied with 3% cysteine in 1/9x MMR, pH 7.8, 10 minutes post fertilization and were then ready for manipulation. Embryos were staged using the Nieuwkoop-Faber developmental table (70). For inhibition of zygotic transcription, 5 pg of α-amanitin (Millipore Sigma) was injected per zygote. Combinatorial inhibition of Otx1/Vegt/Foxh1 was performed as previously described (14) using morpholino antisense oligonucleotides (Gene Tools, LLC) with the sequences 5’-ATGACATCATGCTCAAGGCTGGACA-3’ for *otx1*, 5’-TGTGTTCCTGACAGCAGTTTCTCAT-3’ for *vegt*, and 5’-TCATCCTGAGGCTCCGCCCTCTCTA-3’ for *foxh1*.

### H3K4me1 ChIP-seq and Bioinformatics

ChIP-seq on 1% formaldehyde-fixed dissected *X. tropicalis* embryonic tissues was performed as previously described (9) using an anti-H3K4me1 antibody (Abcam). ChIP-seq libraries were generated using Nextflex ChIP-seq kit (Bioo Scientific), analyzed using an Agilent Bioanalyzer 2100, quantified using KAPA qPCR and sequenced using Illumina instruments at the UC Irvine Genomics High Throughput Facility.

ChIP-seq reads were aligned to the *X. tropicalis* genome v9.0 (71, 72) using Bowtie 2 v2.2.7 (73) with default options. For the rest of the analysis, the set of unique aligned reads for each dataset were down sampled to 15 million reads to enable fair comparison across datasets. To visualize the ChIP-seq tracks, IGVtools functions *sort* (default options) and *count* (-w 25 -e 250) were used to generate the TDF files, which were then loaded into IGV v2.3.20 Genome Browser (74). ChIP-seq peaks were called against stage 10 input DNA (10) using MACS2 v2.0.10 (75) with a p-value cut-off < 10e^−9^ but otherwise default options.

Promoter peaks were defined as peaks < 500 bp from transcription start sites, while enhancer peaks were defined as peaks in intronic and intergenic regions excluding promoter regions. Super-enhancers were identified using the ranking of super-enhancer algorithm (15). H3K4me1-flanked nucleosome free regions (NFRs) were identified using HOMER v4.7 (42) with the function *annotatePeaks*.*pl* –style histone –nfr and a p-value cut-off < 10e^−9^ but otherwise default options. *De novo* motif analysis was performed on 50bp windows from the center of the NFR peaks using DREME (76), and motifs were matched to databases using TOMTOM (77). Genes were assigned to peaks using Bedtools v2.19.1 (78) based on proximity.

### RNA-seq and Bioinformatics

The spatial (ectoderm, mesoderm and endoderm) (17) and temporal (4-9 hours post-fertilization) (18, 37) expression of genes were analyzed using published RNA-seq datasets. Additional RNA-seq datasets were generated whereby RNA was isolated from fifteen individual NF 10.25 (7 hpf) embryos, five each from 3 separate clutches, were subject to Smart-Seq2 cDNA synthesis followed by tagmentation (79) to create libraries. Note that only the genes that do not have maternal expression (TPM > 1 at 0 hours post-fertilization) were considered for the spatial and temporal expression analysis of zygotic genes. Reads were aligned using RSEM v.1.2.12 (80) and Bowtie 2 v2.2.7 (73) to *X. tropicalis* genome v9.0 (71, 72) resulting in expression in transcripts per million. To measure the coefficient of variation (COV) of gene expression, the three time-course datasets were aligned to each other based on reported hours post-fertilization. The COV was calculated as the standard deviation over the mean for each time point per gene. Gene set enrichment analysis was performed using the GSEA software (81, 82) with the pre-ranked option using our ranked genes based on COV.

## Acknowledgements

This work was made possible, in part, through access to the Genomic High-Throughput Facility Shared Resource of the Cancer Center Support Grant (P30CA-062203) at the University of California, Irvine, and NIH shared instrumentation grants 1S10RR025496-01, 1S10OD010794-01, and S10OD021718-01. We thank *Xenbase* for genomic and community resources (http://www.xenbase.org/, RRID: SCR_003280), and the University of California, Irvine High Performance Computing Cluster (https://hpc.oit.uci.edu/) for their valuable resources and helpful staff. This research was funded by the following grants awarded to K.W.Y.C.: National Institute of Health R01 GM126395 and National Science Foundation 1755214.

## Supplemental Information

**Figure S1.**
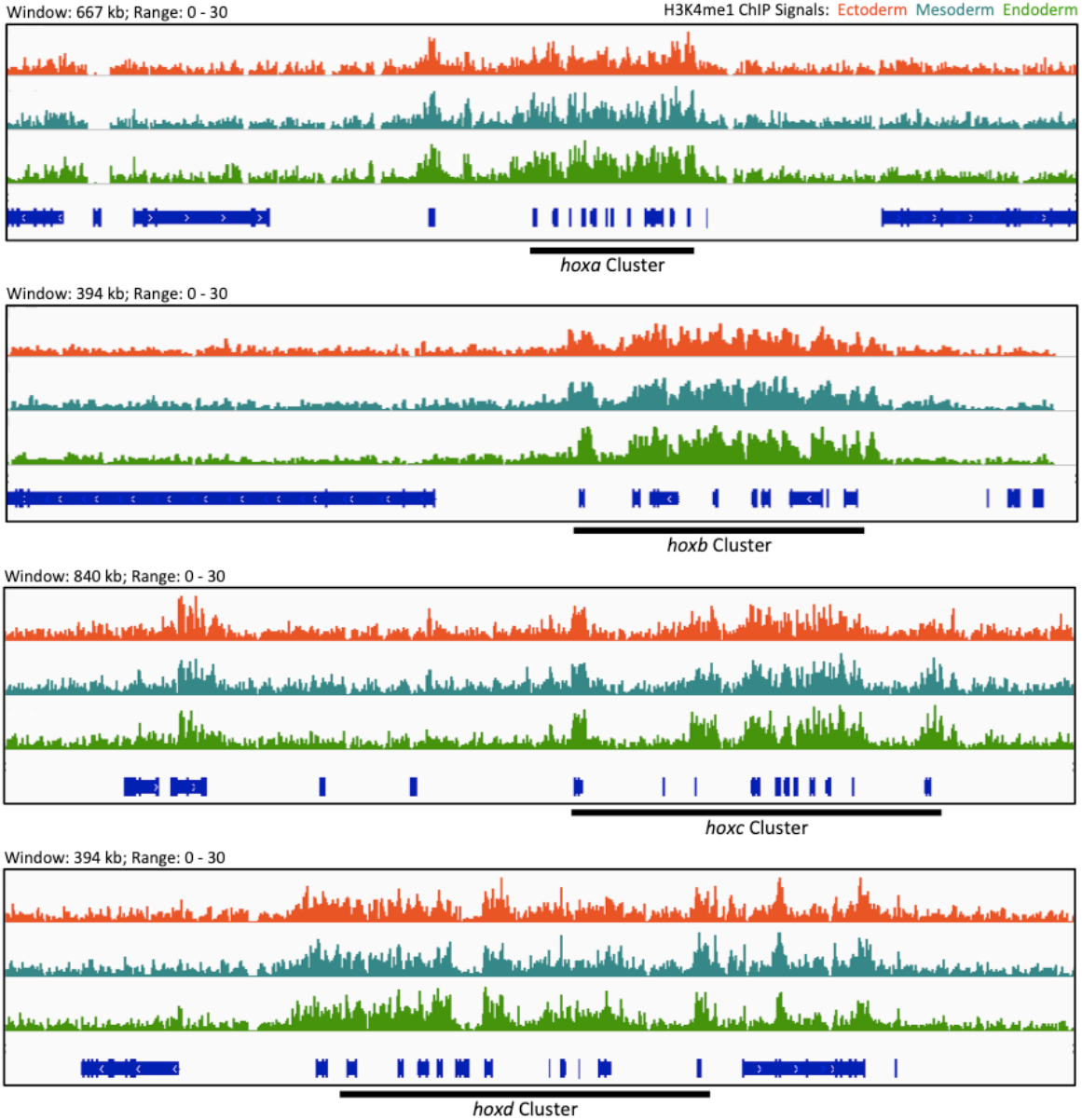
Genome browser view of germ layer-specific H3K4me1 marking in the hoxa, hoxb, hoxc and hoxd clusters loci.

**Figure S2.**
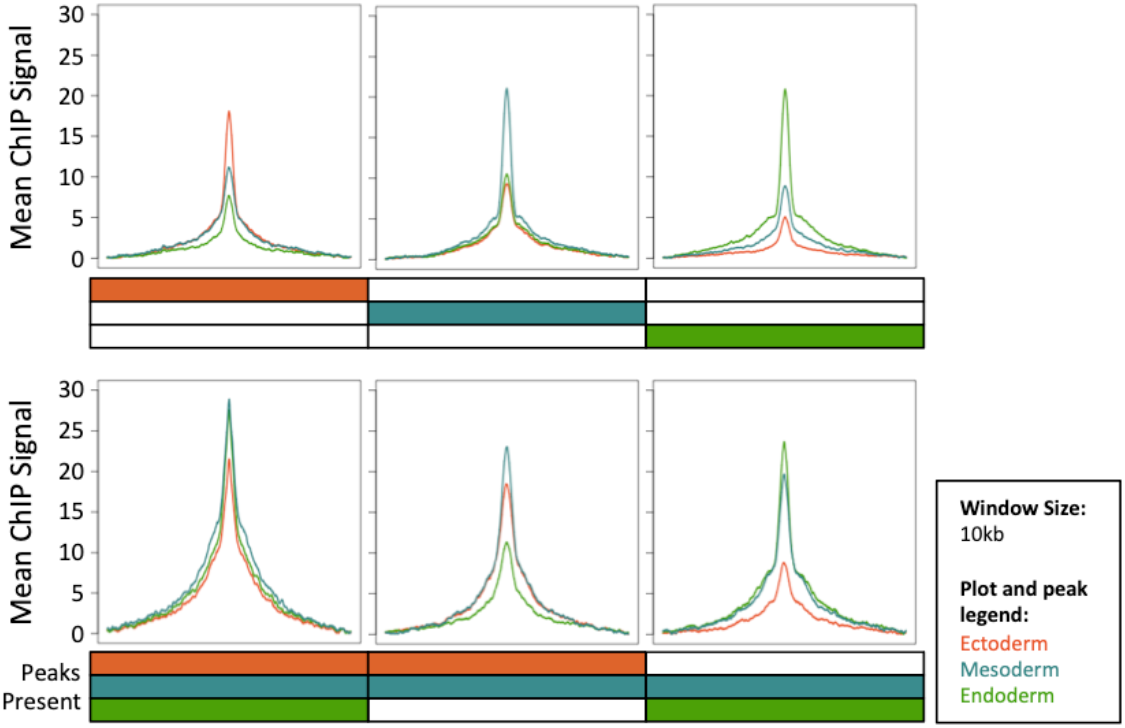
Mean ChIP signal of H3K4me1 ChIP-seq in unique and shared peaks between the three germ layers. Mean ChIP signal is measured as the mean read coverage across the 10 k.b. window for each peak region. For each dataset, the lowest mean ChIP signal is subtracted from the rest of the values within the window such that each plot starts at zero.

**Figure S3.**
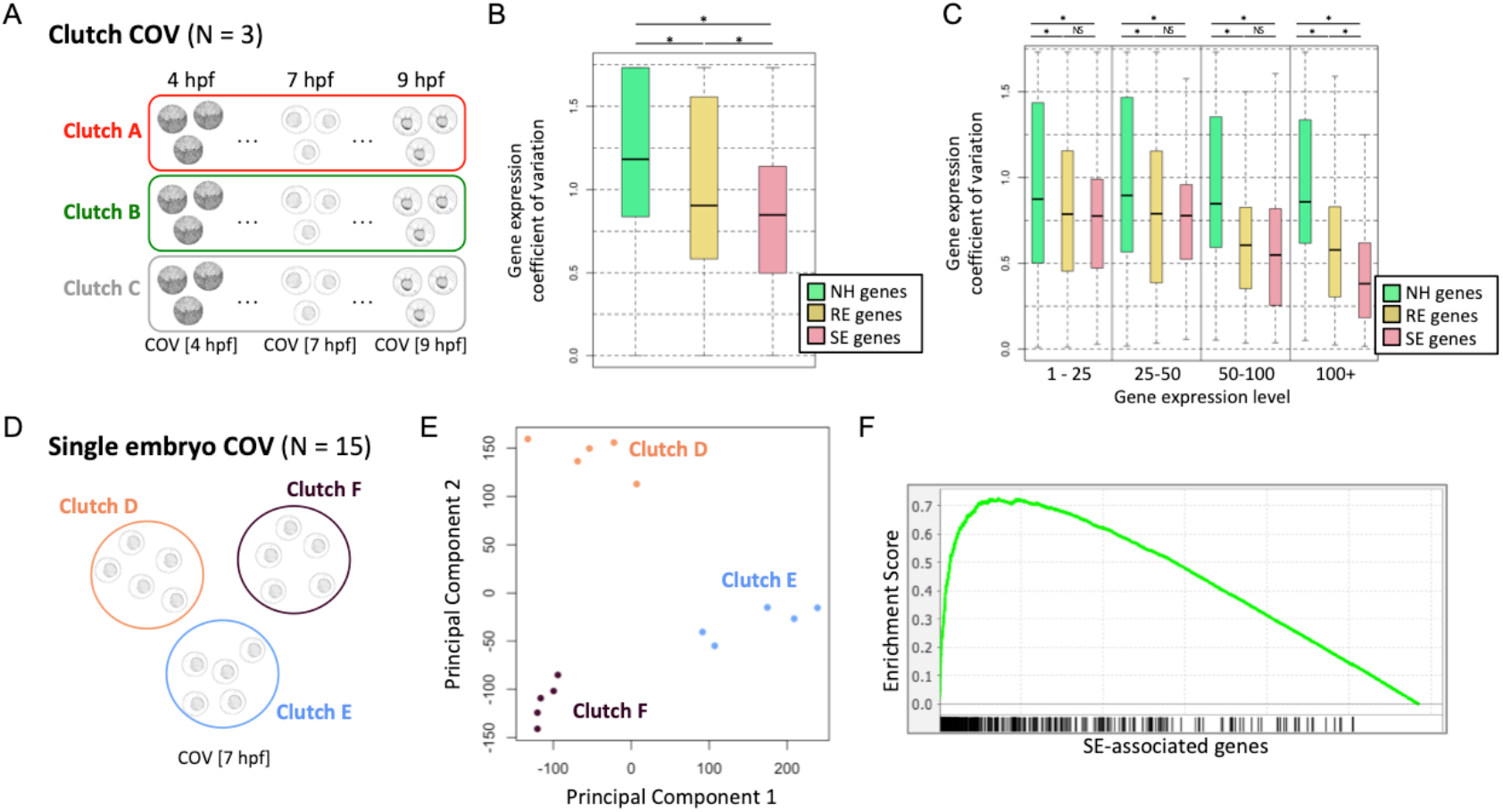
Robustness of gene expression. (A) Measuring the coefficient of variation (COV) between published RNA-seq datasets, which were generated from pooled embryo samples from 4 – 9 hpf with 0.5 hour intervals. The COV is measured for each gene for each time point. (B) COV of gene expression from the RNA-seq of three separate embryo clutches from 4 – 9 hpf. Displayed are the COV for non-H3K4me1 (NH) associated, RE-associated and SE-associated genes. (C) COV of gene expression for genes that are either non-H3K4me1 (NH) associated, RE-associated or SE-associated genes. The expression levels are binned as 1-25, 25-50, 50-100 and 100+ TPM to account for variation caused by the noisy expression at low TPMs. Significance is tested by Wilcoxon Rank-Sum test. * = p-value < 0.01 and NS = not significant. (D) Measuring the COV among 15 embryos from 3 clutches at 7 hpf. (E) Principal component analysis of the transcriptome from the 15 embryo samples. (F) Gene set enrichment analysis of SE-associated genes where genes are ranked based on COV. Each vertical bar at the bottom represents an independent SE-associated gene.

**Figure S4.**
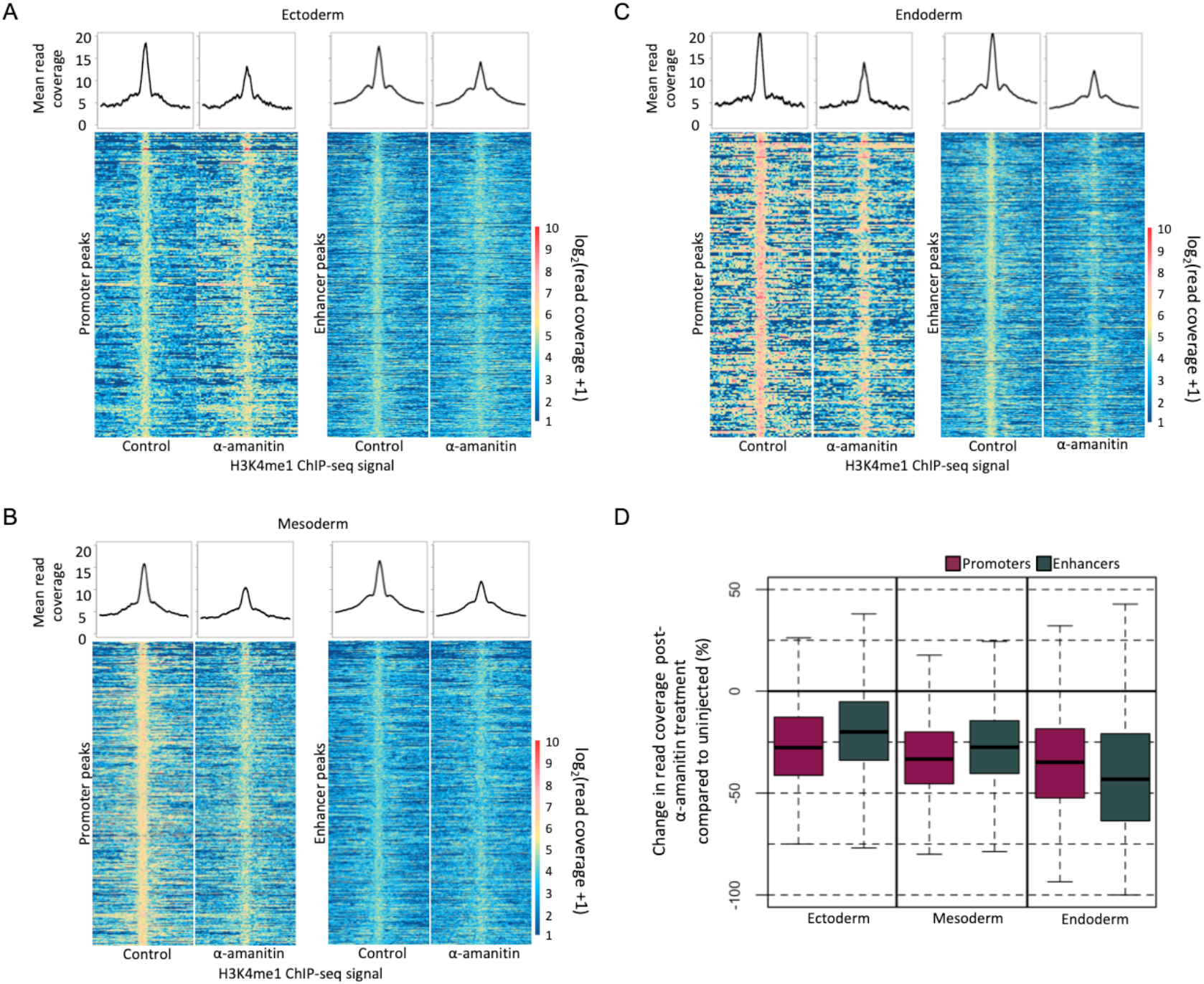
H3K4me1 ChIP-seq signal in control and α-amanitin injected germ layer tissues comparing promoter and enhancer peaks. (A-C) Read coverage of H3K4me1 ChIP-seq of control and α-amanitin injected tissues in promoters and enhancers. The set of displayed promoter and enhancer peaks were identified from the control H3K4me1 ChIP-seq. (D) Quantification of change in read coverage in promoter and enhancer peaks after α-amanitin treatment.

**Figure S5.**
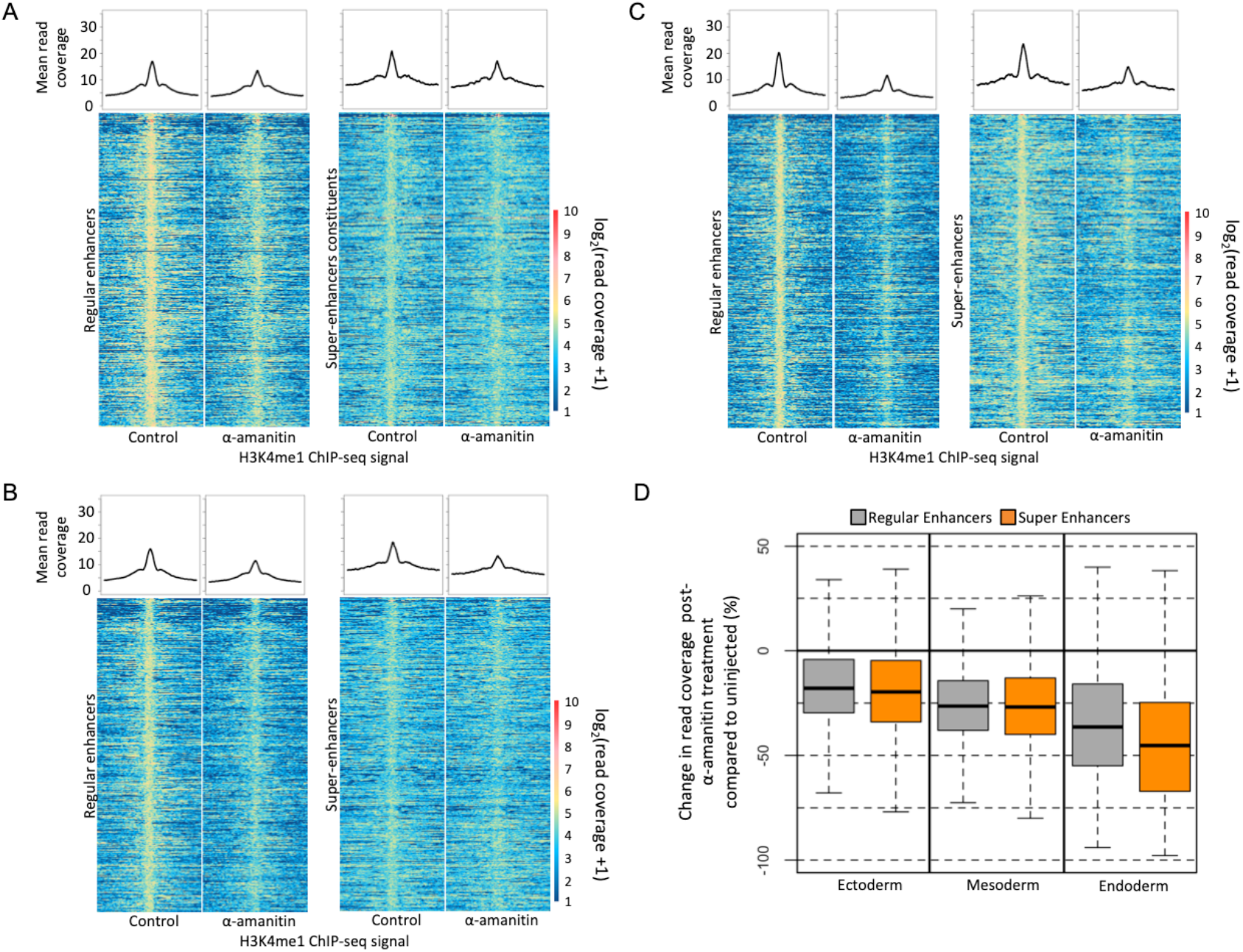
H3K4me1 ChIP-seq signal in control and α-amanitin injected germ layer tissues comparing regular and super-enhancer peaks. (A-C) Read coverage of H3K4me1 ChIP-seq of control and α-amanitin injected tissues in REs and SEs. The set of displayed RE and SE peaks were identified from the control H3K4me1 ChIP-seq. (D) Quantification of change in read coverage in RE and SE peaks after α-amanitin injection.

**Figure S6.**
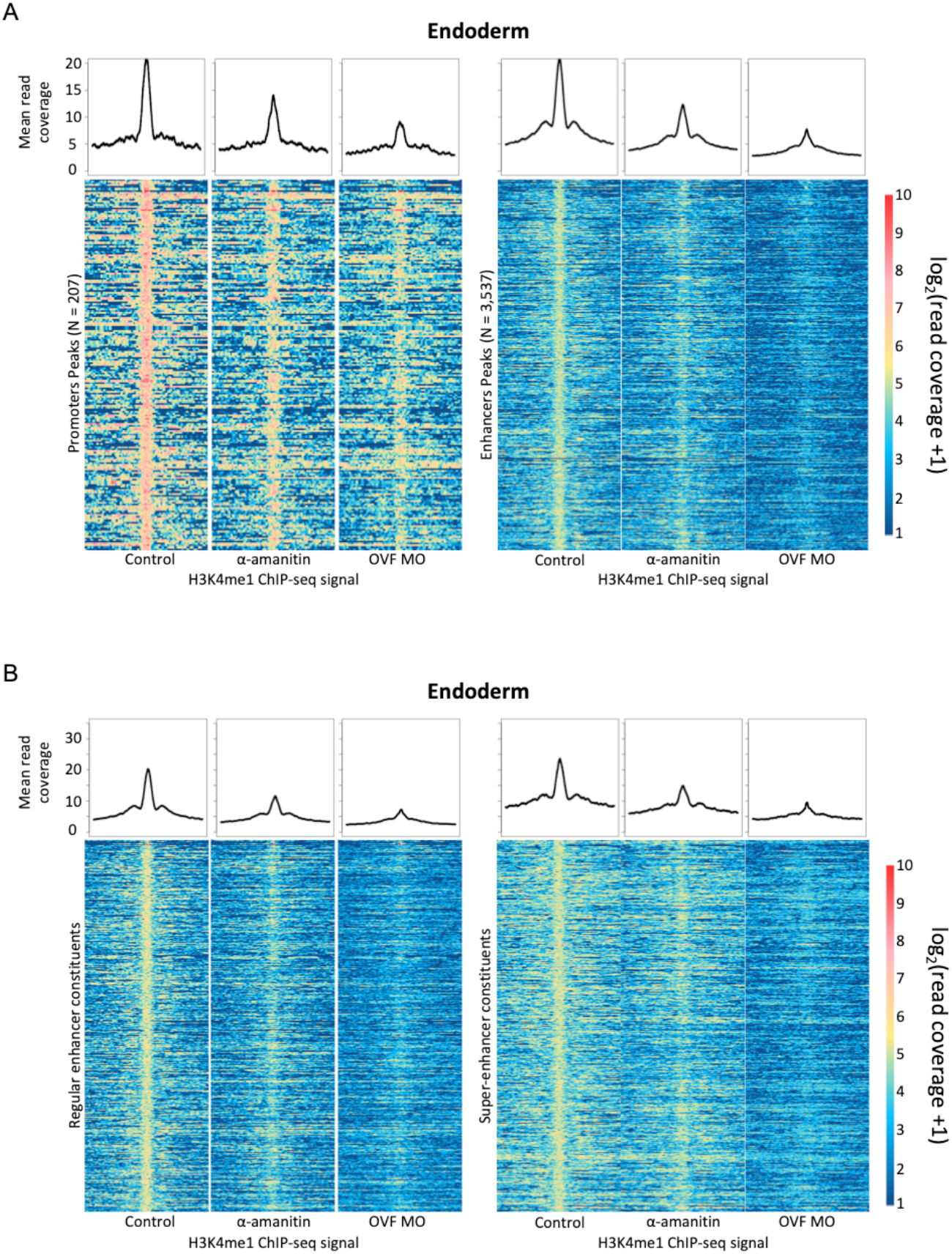
H3K4me1 ChIP-seq signal in control, α-amanitin and OVF MO injected endodermal tissues. (A) Read coverage of H3K4me1 ChIP-seq of control, α-amanitin injected and OVF MO injected endodermal tissues in promoters and enhancers. The set of displayed promoter and enhancer peaks were identified from the control H3K4me1 ChIP-seq peaks. (D-E) Read coverage of H3K4me1 ChIP-seq of control, α-amanitin injected and OVF MO injected endodermal tissues in REs and SEs. The displayed regular and SE peaks were identified from the control endodermal H3K4me1 ChIP-seq peaks.

**Figure S7.**
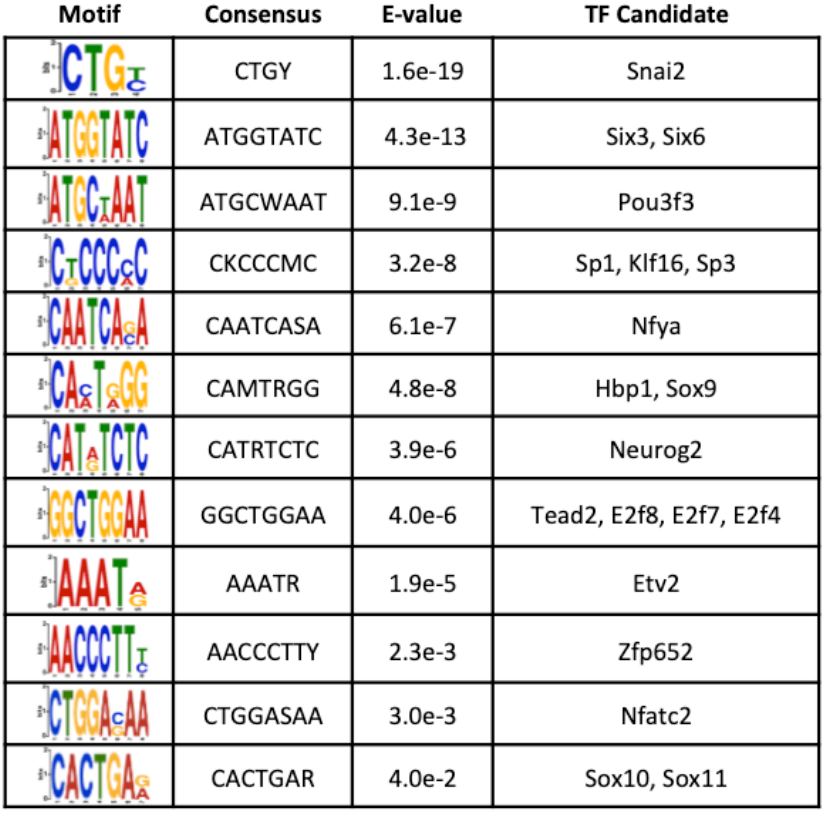
Enriched motifs under H3K4me1-flanked NFR regions of H3K4me1 ChIP-seq that were gained after maternal endodermal TF knockdown compared to control.

## Notes

### Competing Interest Statement

The authors have declared no competing interest.

